# Targeted Delivery of Nucleic Acid and Protein Cargos into Primary Human Hematopoietic Stem Cells Using Bacteriophage T4

**DOI:** 10.64898/2026.01.28.702426

**Authors:** Chelsea E Stamm, Jingen Zhu, Cory Davis-Vargas, Obaiah Dirasantha, Karthika Thankamani, Venigalla B Rao

## Abstract

Here, we report, for the first time, delivery of mRNA and/or protein payloads into primary adult human hematopoietic stem cells (HSCs) using a bacteriophage-derived nanoparticle vector. We have been developing a new category of bacteriophage T4-engineered nanoparticles, termed “artificial viral vectors” (AVVs), for delivering therapeutic nucleic acid and protein complexes into human cells. Using a defined *in vitro* assembly-line platform, we decorated the capsid surface with mRNA and protein complexes through two outer capsid proteins, Hoc (highly antigenic outer capsid protein) and Soc (small outer capsid protein). First, we displayed a Hoc-protein G fusion protein to which HSC-targeted monoclonal antibodies (mAbs) are attached. Then we decorated the nanoparticle with the Soc-fused HIV-TAT molecule to create a positively charged capsid surface with cell penetrating function. Reporter mRNA molecules are then displayed on the capsid surface and the nanoparticle is coated with lipids. Such T4-AVVs transduced HSCs and delivered GFP and Luciferase reporter mRNAs at levels as high as 20% efficiency and exhibited targeted Ab-dependent phenotypes. Furthermore, we demonstrate simultaneous delivery of both protein and mRNA payloads, by decorating the capsid with mRNA and ∼507 kDa tetrameric β-galactosidase. The AVV-transduced HSCs maintain viability, and once optimized, the T4-AVV platform will provide enormous versatility to target various types of human cells and deliver next generation therapies for cancer and genetic diseases.

## Introduction

Curing a genetic disease remains a major global health care challenge. Genetic diseases comprise a vast collection of genome variations affecting various cell types and organ systems and can manifest in a range of symptom severities. Therefore, designing therapies or cures, which is possible for a tiny fraction of genetic diseases, often requires exceedingly costly and time-consuming individualized development. For example, treatments for sickle cell disease and β-thalassemia rely on autologous hematopoietic stem cells (HSCs), expanded, and genetically altered *ex vivo* (1). The patient must undergo chemotherapeutic conditioning to create an available niche before the corrected cells are delivered back to the patient. While successful, significant challenges remain regarding collection and *ex vivo* expansion of HSCs (2), including the up-front financial burden of gene editing, chemotherapy regimen and allogenic transfer (3), and delayed side effects such as the development of leukemias (4). Novel and robust treatments that reduce these obstacles are desperately needed.

Targeted gene therapies delivered *in vivo* would eliminate the need for *ex vivo* editing and expansion and patient conditioning. Indeed, lipid nanoparticles (LNPs) carrying base-editing enzymes were delivered *in vivo* to cure a seven-month-old infant of severe carbamoyl-phosphate synthetase 1 deficiency (5). While this was a significant advancement, much work remains to overcome several technical and biological barriers before *in vivo* gene therapies will be widely accessible. The nanoparticle vehicle must be large enough to carry gene editing machinery including a gene editing enzyme such as Cas9 and gRNA, and donor DNA where homology driven repair (HDR) is also essential. Even monogenetic diseases can be caused by different mutations, requiring customizable and tunable nanoparticles. In addition, the cell type being edited, for example the HSC, comprises a small proportion of the total cells and must be specifically and efficiently targeted. Finally, the efficiency of genome repair in primary cells is variable (6, 7), due to several factors including pre-existing immunity to delivering viral vectors and bacterial editing enzymes (7, 8), innate immune recognition of the donor DNA and differences in nanoparticle uptake, gene expression and preferential repair pathways among varying cell types (6, 9). Thus, being able to deliver the gene editing payload effectively *in vivo* is of utmost importance, and extremely challenging using the current viral or non-viral platforms.

Bacteriophage (phage) T4 (T4) is emerging as an appealing candidate for designing novel and effective gene therapy delivery nanoparticles. Phage T4 is stable at ambient temperature and safe upon intravenous injection (10). Its 120 x 86 nm head (capsid) has a large capacity to encapsidate any linear DNA, or a combination of DNAs, up to 170 kb. In addition, the two surface accessory proteins, the small outer capsid protein (Soc) and the highly antigenic outer capsid protein (Hoc) can tolerate fusion to foreign peptides, or large full-length proteins, without compromising their ability to attach to the capsid. Up to 1025 copies of foreign proteins can be attached to a single capsid using both Soc (870 copies) and Hoc (155 copies) (11). These features allowed us to establish an efficient *in vitro* “assembly-line” platform to create customizable T4 nanoparticles to optimize gene therapy delivery. Indeed, we have used such nanoparticles, referred to as artificial viral vectors (AVVs) to transduce immortalized cell lines such as the human embryonic kidney 293 (HEK293) cells. A variety of genomic remodeling events including large gene delivery and expression, gene silencing and replacement, could be accomplished using the T4-AVVs, greatly expanding the scope of therapies beyond the current adeno associate virus (AAV) and lentivirus (LV) vector models (12).

However, whether the phage-based T4-AVVs can deliver the cargo into primary human cells remained an open question. Transduction into primary cells is essential for clinical applications but generally more challenging when compared to the cell line models (6). Unlike primary cells, the cultured HEK293 cells are more permissible for delivery, having high replication rates and minimal innate immune receptors. Here, as a first step towards targeted *in vivo* delivery, we show successful delivery of both mRNA and protein into primary human HSCs by phage T4-AVVs. By exploiting the modular engineering capacity of the T4 platform, we created nanoparticles decorated with HSC-targeting antibodies and reporter mRNA or protein and coated the particle with lipid. The HSC-targeting antibody positioned at the tip of the ∼18 nm fibrous Hoc fiber likely provided an extended reach to recognize the cognate receptor on the cell surface, while the lipid-coating facilitated entry of the nanoparticle into the HSC. As high as 20% transduction efficiency was achieved using primary HSCs isolated from adult blood. To our knowledge, this is the first report demonstrating phage vector-based transduction of nucleic acid and protein cargos into primary human HSCs, thus providing a proof-of-concept that the large capacity phage T4-AVVs can be engineered for potential stem cell gene therapy applications.

## Results

### Engineering T4-AVV for primary human HSC transduction

We set out to build a T4-AVV that would enter primary HSCs and produce a measurable reporter signal. We modified our previous T4-AVV assembly-line platform (12) starting with the highly acidic phage T4 empty head nanoparticles lacking Hoc and Soc and with each engineered gp23 major capsid protein containing an insertional loop mutation of nine alternating Asp and Glu residues (Figure 1A) (12). We first decorated the nanoparticle with the ∼18 nm long Hoc fiber fused to a 142 amino acid immunoglobulin binding <u>G</u>B domain of Protein <u>G</u> (GG-Hoc). The GG domain binds with high affinity to the Fc domain of all human IgG subclasses (13). We then attached the HSC-specific monoclonal antibodies (mAbs) that can target the nanoparticles to HSCs (Figure 1A, I and III). In addition, we displayed Soc-fused cationic 23-amino acid peptide HIV TAT (14) or the genome editing Cas9 enzyme to the capsid surface in order to capture the GFP or Luciferase reporter mRNA molecules (Figure 1A, II and IV; see below for details). Initially, we chose mRNA as the transduction signal because it can be produced with modified nucleotides to improve stability with reduced immune recognition (15–18). Finally, we coated the negatively charged capsid with cationic lipids (Figure 1A, V). The copy number of each component that is now part of the nanoparticle structure was quantified by SDS-PAGE using internal gp23 capsid protein control (Figure 1B). We routinely achieved 50-100% maximum occupancy of Soc-fused TAT and Hoc-fused-GG-mAb while the occupancy was lower (∼30%) for the much larger ∼170 kDa Cas9-Soc (Figure 1C and 2C, and see below for details).

**Figure 1.**
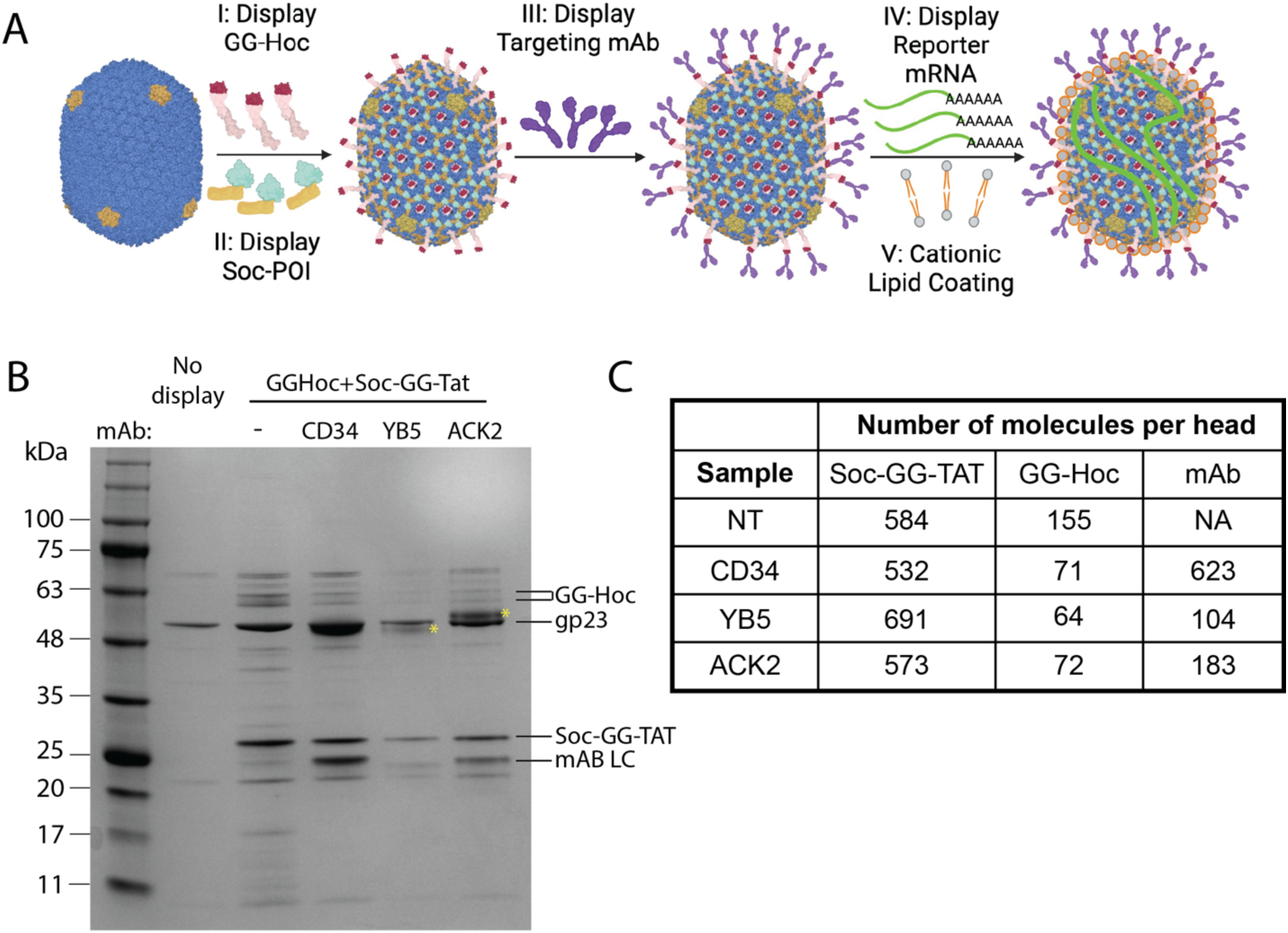
Engineering T4-artificial viral vectors for primary cell transduction. **A**. Schematic detailing the construction of T4-AVV for HSC transduction. Rounds of *in vitro* display are used to coat the T4 capsid (PDB 8GMO) in effector molecules. POI, protein of interest; mAb, monoclonal antibody; mRNA, messenger RNA. **B**. Example of display efficiency analysis. Different T4-AVV preparations were analyzed by SDS-PAGE. CD34, YB5 and ACK denote separate targeting mAb. LC, light chain. Resolved mAb heavy chains are marked with an asterisk. **C**. Quantification of the gel in **B**. The major capsid protein, gp23 is used as an internal standard for copy number.

**Figure 2.**
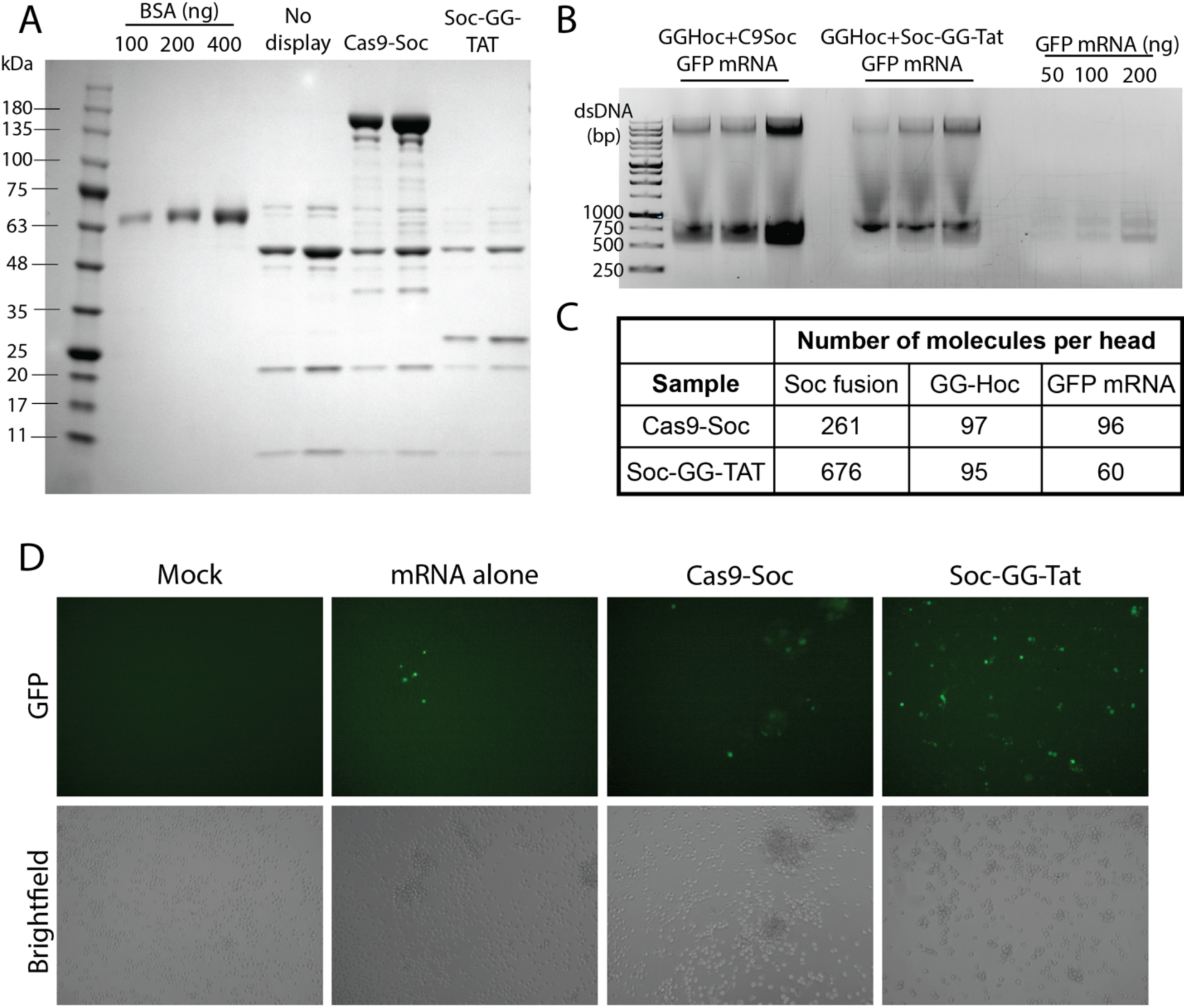
Optimization of mRNA display on T4-AVV. **A**. T4-AVV preparations displaying either Cas9-Soc or Soc-TAT were analyzed by SDS-PAGE. BSA and heads with no display were loaded as quantification controls. **B**. GFP mRNA displayed on T4-AVV via Cas9-Soc or Soc-Tat was visualized by agarose gel electrophoresis. **C**. Quantification of molecules displayed from gels in **A-B**. **D**. Representative fluorescence (top) and brightfield (bottom) microscopy of HSC transduced for 24 h with T4-AVV displaying GFP mRNA via Cas9-Soc or Soc-Tat. Lipofectamine Stem was used to coat the T4-AVV and for the mRNA-LNP control.

### T4-AVVs are not toxic to HSCs

Prior to measuring transduction, we evaluated if T4-AVVs are toxic to HSCs and the level of tolerance. We prepared T4-AVVs with GG-Hoc, a representative display mAb, and lipid, and transduced HSCs. After 48 hours, cells were removed from the well, stained with 0.2% trypan blue and counted using an automated counter. Cell number was not different between mock transduced cells and any T4-AVV treatment (Supplemental Figure 1A). Importantly, viability, as measured by trypan blue exclusion, for all groups was between 95-98% (Supplemental Figure 1A). As an alternative quantification of viability, we prepared T4-AVVs with and without lipid coat or targeting mAb, transduced HSCs at increasing MOIs for 48 hours, and measured luminescence with Cell Titer Glo. The luminescence in cells transduced with T4-AVV lacking a lipid coat remained similar up to MOI 1 × 10^6^. When T4-AVV with and without targeting mAb were coated with lipid we observed a small but dose dependent decrease in luminescence. In all cases, the lowest viability was observed in cells transduced at MOI 5 × 10^6^ but the viability never decreased below mock transduced cells (Supplemental Figure 1B). Lastly, to observe the cell morphology, we stained the cells directly in the well with 0.2% trypan blue for five minutes, washed and visualized immediately under bright field. Mock transduced cells were distributed evenly across the well, and we observed no trypan blue staining (Supplemental Figure 1B). We observed some aggregation in cells treated with lipid-coated T4-AVV but no trypan blue uptake. Finally, we captured rare instances of trypan blue uptake in HSCs treated with mAb antibody coated T4-AVV, consistent with the calculated viability (Supplemental Figure 1C). Taken together these data indicate that T4-AVVs are not toxic to HSCs.

### Engineering T4-AVV to display reporter mRNA

To quantitatively measure transduction of HSCs, we sought to deliver reporter mRNA molecules via T4-AVV. As mRNA cannot be packaged inside the capsid, we used two approaches to engineer mRNA display onto the T4-AVV surface. We previously established that Cas9 fused to Soc and displayed on the capsid could be used to capture and deliver gRNA or an mRNA (12). In addition, we had previously shown that transduction efficiency was improved by decorating the capsid with the Arg-Lys rich HIV peptide TAT to Soc (12, 14). We hypothesized that the negatively charged mRNA molecules would electrostatically bind to capsids displaying Soc-TAT. To compare these mRNA display mechanisms, we used our assembly-line strategy to prepare T4-AVVs coated with GG-Hoc, Cas9-Soc, or Soc-TAT (note that Soc-TAT also contained a GG domain between Soc and TAT because it makes Soc-TAT more soluble in addition to providing sites for Ab binding), and GFP mRNA. The average number of each of these molecules incorporated into the AVV nanoparticle is 96 copies for GG-Hoc, 676 copies for Soc-TAT, and 261 copies for Cas9-Soc (Figure 2A and C). T4-AVVs displaying Cas9-Soc bound on average 96 copies of GFP mRNA whereas 60 copies were bound to Soc-TAT displaying AVVs (Figure 2B-C).

To determine if either of these GFP mRNA-AVVs could transduce HSCs, we prepared Cas9-Soc or Soc-TAT coated T4-AVVs displayed with GFP mRNA, transduced cells for 24 hours, and compared the GFP signal to a positive control in which cells were transfected directly with mRNA-lipid mixture (referred to as “mRNA-LNPs”; see Methods). Excitingly, we observed fluorescent cells in HSCs transduced with T4-AVVs and there were more fluorescent cells per field in both the T4-AVV transduced cells compared to the control mRNA-LNP transfection (Figure 2D). The fluorescence in cells transduced with Cas9-Soc coated AVVs was less intense than the Soc-TAT coated AVVs and the cells had aggregated. In contrast, individual cells were transduced with the GFP signal in Soc-TAT coated T4-AVVs and the brightness was similar to that of the positive control cells directly transfected with mRNA-LNPs (Figure 2D). Due to the higher efficiency and fluorescence phenotype, we chose Soc-TAT AVVs to deliver mRNA for further studies.

### Targeting HSC surface molecules enhances transduction by T4-AVVs

Primary HSCs can be characterized by their surface proteins. The glycoprotein CD34 is the most commonly used marker to isolate the HSC population from whole blood. In addition, the receptor tyrosine kinase c-KIT or CD117, while not unique to HSCs, is also highly expressed on their surface. We hypothesized that displaying a mAb against HSC-enriched surface proteins on T4-AVVs would increase nanoparticle binding and/or internalization and improve transduction efficiency. To test this, we prepared T4-AVVs displaying luciferase mRNA and coated with mAbs targeting CD34 or CD117, a nonspecific control Ab (goat α-IgG1), or no mAb, transduced the HSCs for 24 hours and measured luminescence. The luminescence in cells transduced with non-targeting T4-AVV was nearly two logs higher than mock transduced cells and coating with a nonspecific mAb did not improve the signal (Figure 3A). In contrast, luminescence increased when cells were transduced with T4-AVVs displaying mAb against CD34 or two different mAbs against CD117 (YB5 and ACK2, Figure 3A). In all cases, transduction with T4-AVVs produced higher luminescence than the positive control mRNA-LNP transfection (Figure 3A).

**Figure 3.**
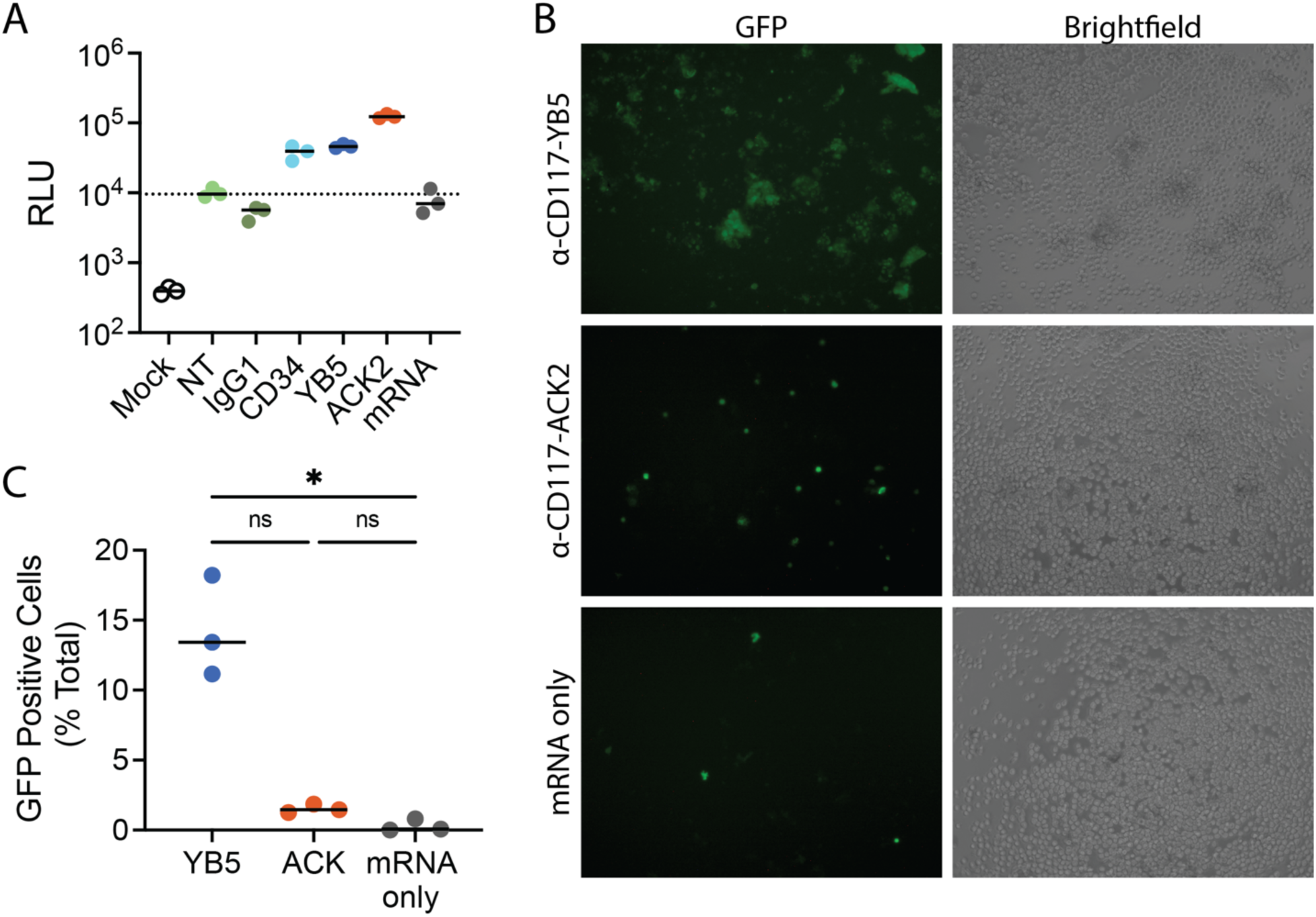
Targeting HSC surface molecules improves transduction by T4-AVV. **A**. Relative luminescence units (RLU) in HSC mock transduced or transduced for 24 h with T4-AVV displaying luciferase mRNA and the indicated targeting mAb. NT, no targeting. Samples were performed in biological triplicate. **B**. Representative fluorescent and bright-field images of HSC 24 hpt with T4-AVV displaying GFP mRNA and different α-CD117 mAb or directly transfected with GFP-mRNA. **C**. Quantification of samples in **B**. Three fields of at least 100 cells were counted for each condition. Data in **A-C** are representative of three independent experiments.

Interestingly, we observed a difference in the luminescence signal between the two CD117 targeting mAbs, with the ACK2 mAb showing 2.5-fold higher luminescence than the YB5 mAb. To validate this difference, we repeated the targeting experiment by preparing T4-AVVs displaying GFP mRNA and coated with the CD117 mAbs, YB5 or ACK2, and visualized the cells under fluorescent microscopy. Cells transduced with the YB5 T4-AVVs aggregated and produced fainter fluorescence across the clumps (Figure 3B). In contrast, cells transduced with ACK2 T4-AVVs remained separated, and were brighter but fewer cells per field were transduced (Figure 3B-C). In both cases the transduction efficiency as measured by the percentage of GFP positive cells, was 18-160 times higher than non-targeting GFP mRNA transduction (Figure 3C), consistent with the luminescence data. Taken together we concluded that targeted AVVs enhanced transduction efficiency and CD117 is the preferred target.

### The type of lipid coat affects HSC transduction

Previously, T4-AVVs coated with the cationic Lipofectamine 2000 gave high levels of transduction into HEK293T cells. In order to achieve the highest transduction while maintaining HSC viability, we used lipid coats that were reported to be optimal for primary cell and mRNA transduction. CD117-targeted, GFP mRNA-displaying T4-AVVs were coated with either Lipofectamine Stem (Stem), Lipofectamine MessengerMAX (MM) or LipofectamineCRISPRMAX (CM), and tested for their transduction efficiency into HSCs after 24 hours (Figure 4A). In addition, we have also observed the cells for any changes in cell morphology (Figure 4B). As before, cells transduced with YB5 mAb-targeted T4-AVVs showed clumping behavior (Figure 4B), but the number of fluorescent cells ranged from 8-20% with CM showing the best transduction efficiency (Figure 4A). Similarly, cells transduced with ACK2-targeted CM-coated T4-AVVs also showed as high as 6% GFP positivity, and again, consistent with the previous results, the cells remained separated and no clumping was observed. The control mRNA transfections using the respective lipid reagent consistently showed lower efficiencies than with the T4-AVVs (Figure 4A-B).

**Figure 4.**
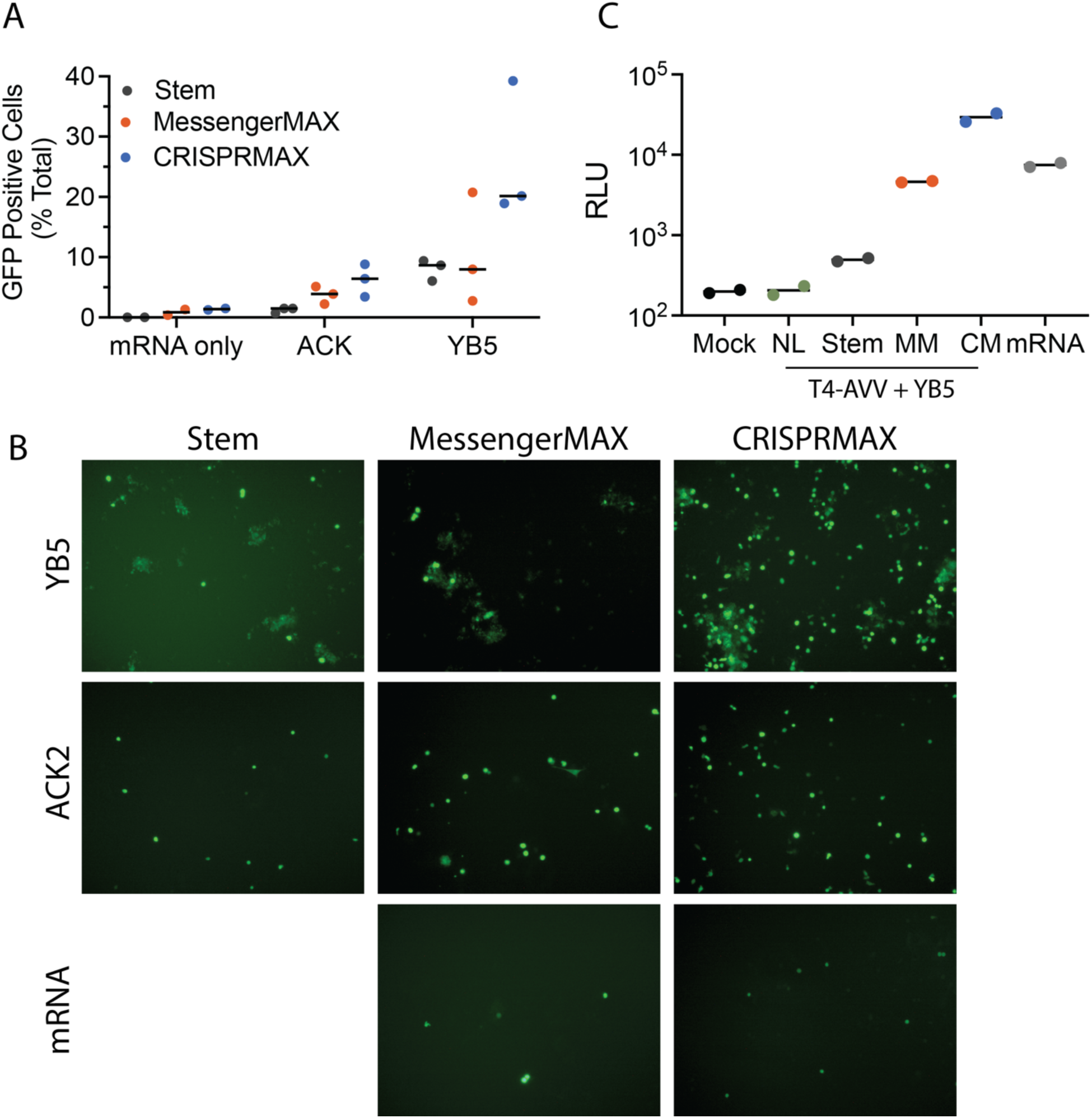
The T4-AVV lipid coating affects HSC transduction. **A.** Quantification of GFP positivity in HSCs 24 hpt with T4-AVVs displaying GFP mRNA, the indicated CD117 mAb and Lipofectamine mixture. Two (mRNA) or three (YB5, ACK2) fields of at least 100 cells were counted for each condition. **B.** Representative fluorescent images from **A**. No cells were transfected by mRNA-LNPs when Stem was used as the lipid coat. **C.** Relative luminescence units (RLU) in HSC 24 hpt with T4-AVVs displaying luciferase mRNA, mAb YB5 and the indicated Lipofectamine mixture. NL, no lipid; MM, messengerMAX; CM, CRISPRMAX. Luminescence was measured in biological duplicates. Data in **A-C** are representative of two independent experiments.

Next, to validate the observed differences, we repeated the lipid transduction experiments using YB5 targeted T4-AVVs displaying luciferase mRNA and measured luminescence 24 hours post-transduction (hpt), (Figure 4C). Mock transduced cells or cells transduced with T4-AVVs without lipid coating as expected only produced low background luminescence. In contrast, we observed luminescence signal in all samples transduced with T4-AVVs coated with lipid, with CM coating again providing the highest efficiency, ∼100 fold higher than the mock control, and ∼5-fold higher than mRNA-LNP transfection control (Figure 4C).

From the above results, we determined that the lipid coating can enhance transduction, to as high as 20% efficiency with CRISPRMAX, leaving substantial room for improvement through screening of various lipids and lipid compositions.

### Delivery of a reporter protein complex into HSCs by T4-AVVs

Delivery of protein molecules, such as gene editing machinery, will be essential for development of future gene therapies. To test if T4-AVV could deliver functional proteins into primary HSCs, we produced Soc-fused β-galactosidase (Soc-β-gal, ∼507 kDa), a tetrameric enzyme that hydrolyzes the β-galactoside bonds in oligosaccharides. We prepared YB5-targeted T4-AVVs coated with Soc-β-gal, transduced HSCs as above, and measured β-galactosidase activity 5 and 24 hpt using the colorimetric substrate o-Nitro-β-D-galactopyranoside (ONPG). We observed increasing ONPG conversion with increasing T4-AVV MOI at both 5 and 24 hpt (Figures 5A-B). Transduction with as little as 5 × 10^5^ MOI produced the β-gal signal above the mock transduced or the non-targeting controls at both time points (Figures 5A-B). The Soc-β-gal activity at the highest dose (5×10^6^ MOI) was higher than the signal from control purified protein (13 ng of Soc-β-gal) at 5 hpt (Figure 5A) and remained similar to the control at 24 hpt (Figure 5B).

**Figure 5.**
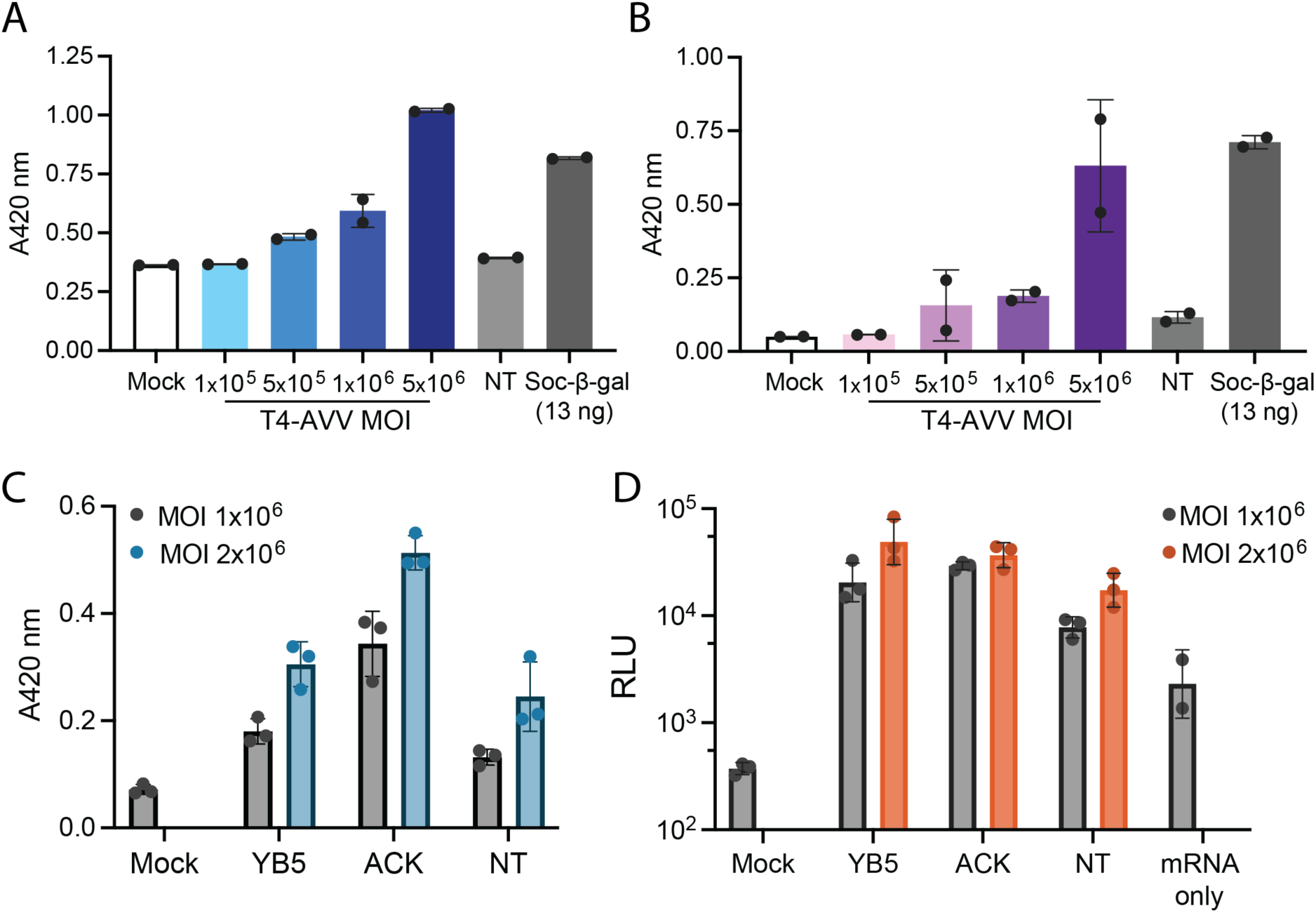
Protein delivery into HSCs by T4-AVVs. **A-B**. Soc-β-gal activity (absorbance at 420 nm) in HSCs 5 hpt (**A**) and 24 hpt (**B**) with increasing doses of T4-AVVs coated in Soc-β-gal. NT, no targeting mAb; MOI, multiplicity of infection. Purified Soc-β-gal was used a positive control for the assay. Samples were measured in biological duplicate. **C-D.** Soc-β-gal activity (**C**) and luminescence (**D**) in HSCs 5 hpt with T4-AVVs coated in a 1:1 molar ratio of Soc-β-gal and Soc-Tat, the indicated mAb and luciferase mRNA. NT, no targeting mAb. Samples were measured in biological triplicate. Data in **A-D** are representative of two independent experiments.

Future gene therapies would rely on simultaneous delivery of both nucleic acids and proteins for gene editing and homology driven repairs. Thus, we set out to determine if T4-AVV could deliver and express both Soc-β-gal and reporter mRNA incorporated into the same nanoparticle. We prepared either YB5- or ACK2-targeting T4-AVVs decorated with both Soc-β-gal and luciferase mRNA. Prior to displaying mRNA, we first incubated the T4 heads with both Soc-β-gal and Soc-TAT at 1:1 molar ratio to display both on the same capsid. The copy numbers of the final T4-AVVs are 65 copies of Soc-β-gal, and 720 copies of Soc-TAT. We then transduced these nanoparticles at two intermediate MOIs (1 × 10^6^ and 2 × 10^6^) into HSCs and measured Soc-β-gal activity and luminescence from the same lysates. Again, there was a dose dependent increase in ONPG conversion in cells transduced with all the T4-AVV constructs (Figure 5C). Furthermore, targeting with YB5 or ACK2 enhanced Soc-β-gal delivery compared to the non-targeting control (Figure 5C). Importantly, the T4-AVVs were also able to deliver luciferase mRNA into the same cells as measured by luminescence. We have also observed a similar dose dependent increase in luminescence for YB5-targeted and non-targeting T4-AVV transduction, and all T4-AVV transductions resulted in greater luminescence than the control cells transfected with luciferase mRNA alone (Figure 5D).

Taken together, the above sets of data show that T4-AVVs not only can deliver large oligomeric protein complexes into primary HSCs but can also do so with both proteins and nucleic acids at the same time when incorporated into the same nanoparticle.

## Discussion

The development of an all-in-one vehicle for *in vivo* gene therapy administration would be invaluable. Here we show, to our knowledge, the first record of primary human cell transduction by bacteriophage T4. We utilized the vast engineering space of our T4-AVV “plug and play” platform to systematically develop and optimize nanoparticles capable of delivering mRNA and proteins into HSCs with no significant effect on cell viability. By establishing this baseline of transduction into primary stem cells, we can further explore the strengths of the T4-AVV platform as a next-generation biological therapeutic.

A unique advantage of the T4-AVV platform is the ability to display biomolecules ranging from small peptides and domains, such as TAT and GG, to large and even multimeric proteins like Cas9 and β-gal as fusions to the T4 surface proteins Hoc and Soc. The incorporation of these effector proteins into the AVV nanoparticle expanded the engineering capacity, further promoting the addition of features including targeting mAbs and mRNA. We took advantage of this flexible engineering space and the sequential assembly-line mechanism for preparing T4-AVVs to perform rapid experimental variation to identify nanoparticles capable of transducing HSCs. By systematically sampling therapeutic molecule combinations, we reached up to 20% transduction efficiency, illustrating the strength and adaptability of the AVV platform. This quick programmability will be an important feature for developing novel T4-based therapeutics.

Safety is a fundamental concern for any biological-based therapeutic. We tested the HSC toxicity of T4-AVVs at increasing MOIs and with various surface biomolecules. Importantly, T4-AVVs were well tolerated as measured by population level viability assays and single cell staining and morphology analysis. Gene therapy schemes that edit stem cells rely on the pluripotency and high replicative capacity of this small cell population (19). The transduction experiments we conducted here lasted 5 to 48 hours, presenting little time for cell expansion. Therefore, the self-renewal and differentiation potential of HSCs following T4-AVV transduction remains an open area of investigation.

Designing a gene editing-based therapy requires a multivalent nanoparticle that can deliver gene editing enzymes, small guide RNAs and HDR DNA donors (8, 20–22). Large fusion proteins can be attached to Soc at either terminus while retaining capsid binding (12, 23, 24). Although the occupancy of large proteins, such as the ∼507 kDa Soc-β-gal, is lower due to their size (65 molecules per head), that these could be delivered at all into HSCs is significant and the signal is robust. In addition, using the modular platform, we were able to display two different Soc fusion proteins on the same capsid and measured delivery of both protein and mRNA simultaneously without compromising either reporter signal. In vaccine design, providing the antigen on the surface and as an expressible DNA increased potency (10, 25). In this case, delivering functional editing enzymes on the nanoparticle surface in conjunction with packaging an expressible DNA may improve editing efficiency and reserves space for co-packaging HDR donor DNA inside the capsid. As such, detecting DNA delivery, gene expression and genome editing in primary cells is the next stage of T4-AVV development and currently under investigation.

A major challenge for *in vivo* gene therapy is delivering the therapeutics to the cell type of interest (7, 8, 26, 27). The T4 Hoc fiber, which occupies 155 binding sites on the capsid extends ∼18 nm from the capsid surface and is well poised to provide targeting functionality (28, 29). Indeed, we incorporated mAb binding into the AVV nanoparticle by displaying Hoc fused to the Fc-binding GG peptide at the tip of the fiber. The Ab binding occurs at the tip through the Fc domain, thus exposing the epitope binding Fab domains at >20 nm from the capsid surface providing considerable reach to contact receptors on the cell surface. The largest improvement in HSC transduction was observed with AVVs coated with α-CD117. Interestingly, we observed differences in cell morphology and transduction efficiency between mAb against two CD117 clones. The YB5.B8 (YB5) clone is against human CD117 and is isotype IgG1, kappa (30). In contrast the ACK2 clone is against mouse CD117, with reports of cross reactivity for human CD117 (31, 32). The source of the antigen may explain the differences in both morphology and transduction efficiency. The human antibody YB5 consistently yielded the most transduction efficiency, but the high affinity for CD117 combined with the icosahedral structure of the AVV nanoparticle may promote cell aggregation. In contrast the cells remained separated when transduced with T4-AAVs coated in the mouse antibody ACK2. Although we observed lower transduction efficiency of GFP mRNA, the cells were brighter and this was reflected in higher luminescence at the population level. These results reflect the difficulty of balancing mAb affinity for efficient cell targeting with maintaining homeostatic cell function.

Once targeted, delivering the large therapeutic nanoparticles across lipid bilayer of primary cells is the next challenge. Cationic lipid-based formulas are effective at delivering nucleic acids and proteins across the anionic lipid bilayer (33, 34). Indeed, in the absence of a lipid coating the T4-AVVs transduced HSCs poorly. As we had previously engineered the anionic capsid for cationic lipid binding (12), we screened for lipid components that could enhance HSC transduction. T4-AVVs coated with Lipofectamine CRISPRMAX provided the highest transduction efficiency. In fact, we observed a 10% increase in GFP positivity and a nearly two log increase in luminescence between cells transduced with CM and the stem cell-specific formulation Lipofectamine Stem, highlighting the importance of identifying the appropriate lipid composition for each cell type. However, traditional cationic transfection lipids carry limitations in moving to *in vivo* delivery including little control over complexation and rapid clearance from circulation (35, 36). Instead, LNPs provide the most promising mechanism for *in vivo* delivery. Durable and safe LNP formulations have been established for mRNA vaccine delivery (37, 38) and gene therapy delivery (5). Although accumulation in the liver is most common (26, 39, 40), modification of the formulation to target other organ systems has been reported (41, 42). Formulating LNPs for T4-AVV delivery will require careful optimization as the AVV nanoparticle is much larger and precisely structured with fixed dimensions than the typical nucleic acid cargos.

In conclusion, we have developed the T4-AVV platform for primary human HSC transduction, marking a significant advancement in bacteriophage-based therapeutic nanoparticle design. How T4-AVVs enter HSCs and are trafficked to deliver cargo appropriately remains unresolved but we established that incorporating targeting moieties into T4-AVVs and modulating the lipid coat provided significant transduction enhancement with up to 20% efficiency. Thus, we have only just explored a fraction of the transduction capacity and experiments to improve the nanoparticle targeting using nanobodies and internalization via novel LNPs are ongoing with the goal of delivering DNA cargo and editing of the genome. Although this work focused on HSCs, the advantages presented here including the nanoparticle safety, modular design, expanded engineering space, and all-in-one delivery could emerge as powerful tools for designing therapies for other diseases where genetic alteration can lead to a cure, such as augmenting the immune system for cancer therapy or rendering cells HIV resistant through receptor deletion.

## Methods

### T4-AVV coating reagents

The targeting antibodies used were α-CD34, clone 581 (Stemcell Technologies, #60013), α-CD117, clone YB5.B8 (ThermoFisher Scientific #14-1179-82), and α-CD117, clone ACK2 (ThermoFisher Scientific #14-1172-82) and the nonspecific antibody anti IgG1 (Abcam #ab98693). The following cationic lipid reagents were purchased from ThermoFisher Scientific: Lipofectamine Stem (#STEM00008), Lipofectamine MessengerMAX (#LMRNA008), Lipofectamine CRISPRMAX (#CMAX00015).

### Recombinant protein expression and purification

Sequences of recombinant proteins are listed in Supplemental Note 1. The genes were cloned into pET28b, transformed into BL21 (DE3) RIPL cells (Agilent #230280) and grown in 1 L Moore’s medium (20 g tryptone, 15 g yeast extract, 8 g NaCl, 2 g dextrose, 2 g Na2HPO4, and 1 g KH2PO4 dissolved in 1 L of Milli-Q water) supplemented with 50 µg/mL kanamycin and 34 µg/mL chloramphenicol to a density of 2-4 × 10^8^ CFU/mL. Protein expression was induced with 0.5 mM isopropyl β-d-1-thiogalactopyranoside (IPTG) for 3-4 hours at 28°C. Cells were harvested by centrifugation and cell pellets were stored overnight at −80°C. All purification steps occur on ice unless otherwise noted. Cell pellets were thawed and resuspended in 40 mL lysis buffer (50 mM Tris, pH 7.5, 300 mM NaCl, 20 mM imidazole) supplemented with EDTA-free protease inhibitor cocktail (Roche, #11836170001), lysed by French press and centrifuged at 34,000 x g for 30 minutes at 4°C to clear debris. The supernatant containing soluble protein was saved, filtered by a 0.22 µm PES filter and applied to a pre-equilibrated HisTrap HP nickel column (Cytiva #17524701) via the ÄKTAStart chromatography system. The column was washed in 30-40 mL wash buffer (50 mM Tris, pH 7.5, 300 mM NaCl, 30 mM imidazole) before the His-tagged protein was eluted on a linear gradient from 30-500 mM imidazole. The fractions containing the protein of interest were pooled, buffer exchanged into gel filtration (GF) buffer (20 mM Tris-HCl and 100 mM NaCl, pH 7.5) and further purified by size exclusion chromatography on a Hi-Load 16/60 Superdex-200 (prep-grade) gel filtration column (GE Healthcare) in GF buffer according to the manufacturer’s instructions. Peak fractions were combined and concentrated using Amicon Ultra centrifugal filter units with 10 kDa molecular weight cut off (Milipore Sigma #UFC901008), quantified by SDS-PAGE against a bovine serum albumin (BSA) standard and stored at −80°C.

### In vitro synthesis of EGFP and Luciferase mRNA

EGFP and luciferase mRNAs were synthesized by *in vitro* transcription using the mMESSAGE mMACHINE™ T7 mRNA Kit with CleanCap™ Reagent AG (ThermoFisher Scientific #A57620-25). A linearized pMK-HA-emGFP template containing a T7 promoter with an AG initiation sequence and flanking 5′ and 3′ UTRs was used for EGFP mRNA synthesis. For luciferase mRNA, the emGFP coding sequence was replaced with a codon-optimized luciferase gene (Supplemental note 1). The transcription templates were generated by polymerase chain reaction using given primers (Forward: 5’ - AGTAATACGA CTCACTATAA GGAGA - 3’, Reverse: 5’ - TTTTTTTTTT TTTTTTTTTT TTTTTTTTTT TTTTTTTTTT TTTTTTTTTT TTTTTTTTTT TTTTTTTTTT TTTTTTTTTT TTTTTGCCGC CCACTCAGAC - 3’). According to the manufacturer’s protocol, *in vitro* transcription reactions (20 µL) contained reaction buffer, ATP, CTP, GTP, and UTP (7.5 mM each), CleanCap™ Reagent AG (6 mM), linear DNA template (25 ng/µL), RNase inhibitor, T7 enzyme mix, and nuclease-free water, and were incubated at 37°C for 3 h. Reactions were treated with DNase I for 15 min at 37°C and purified by LiCl precipitation. Purified mRNA was resuspended in nuclease-free water, quantified by NanoDrop spectrophotometry, and integrity was confirmed by agarose gel electrophoresis.

### T4 heads purification

The 9DE-gp23 10-amber 13-amber hoc-del soc-del ipII-del T4 phage was propagated via *E. coli* B40 (sup^1^, with amber-suppressor). For T4 head production, *E. coli* P301 cells (sup-, no amber suppressor) were infected at an MOI of 5 and incubated at 37°C, 250 rpm for 7 minutes, then superinfected at an MOI of 5 and incubated with shaking for an additional 25 minutes. Cells were harvested by centrifugation (25,000 × *g*, 40 min) and resuspended in 35 mL Tris-Mg buffer (50 mM Tris-HCl, pH 7.5; 5 mM MgCl₂) supplemented with one cOmplete™ mini protease inhibitor cocktail tablet (Millipore Sigma #11836170001) and Benzonase (5 µL; VWR # EM70664-3). Chloroform (1 mL) was added to lyse cells, and the resuspension was shake-incubated at 37°C for 1 h with vigorous pipetting at 30 min. To help separate T4 capsids from cellular debris, 3 mL of 5M NaCl was added, followed by sequential low speed (4000 x *g* for 20 min) and high-speed centrifugation (25,000 x *g* for 1 hour). High speed pellets were soaked overnight at 4°C in 750μL low salt Tris·Mg buffer (50mM pH 7.5 Tris-HCL, 50mM NaCl, 5mM MgCl_2_). The following morning the pellets were resuspended, followed by a low-speed centrifugation (4000 x *g* for 20 min) and a CsCl density gradient centrifugation of the supernatant. Extracted heads were then dialyzed in 2-h intervals against high salt Tris·Mg buffer (50mM pH 7.5 Tris-HCl, 200mM NaCl, 5mMMgCl_2_) then low salt Tris·Mg buffer (50mM pH 7.5 Tris-HCl, 50mM NaCl, 5mMMgCl_2_). Dialyzed heads were further purified by anion exchange chromatography via DEAE-Sepharose or quaternary-ammonium (Q)-Sepharose. Elution fractions containing heads were buffer-exchanged into low salt Tris·Mg buffer (50mM pH 7.5 Tris-HCL, 50mM NaCl, 5mM MgCl_2_) by centrifugal filtration, concentrated, and stored at −80°C. The quantity of capsid particles was determined by SDS-PAGE analysis of the major capsid protein gp23 (48.7 kDa, ∼.75μg gp23 per 10^10^ T4 heads) compared with known quantities of T4 heads or SARS-CoV-2 nucleocapsid protein (Sino Biological).

### HSC culturing and expansion

Media components included DMEM High glucose supplemented with pyruvate and GlutaMAX (ThermoFisher Scientific #10569044), fetal bovine serum (FBS, Avantor Sciences #89510-186), Penicillin-Streptomycin (PS, ThermoFisher Scientific #15140122), StemSpan SFEM II (Stemcell Technologies #9605), StemSpan Human CD34+ expansion supplement (Stemcell Technologies #2691), and UM729 (Stemcell Technologies #72332). Human CD34+ HSCs from cord blood were purchased frozen from Stemcell Technologies (#70008.3, mixed donors) and human CD34+ HSCs from adult mobilized bone marrow were purchased frozen from Seattle Blood Works. To expand, cells were thawed at 37°C and resuspended in DMEM supplemented with 10% FBS and PS, centrifuged at 300 x g for 10 minutes, resuspended in expansion media (Stemspan SFEM II supplemented with 1X CD34 expansion supplement and 1 µM UM729), counted and plated in 24-well plates at a density of 10,000 cells per well. Cells were incubated at 37°C, 5% CO_2_ for 4-5 days, removed from the well with 0.25% Trypsin (ThermoFisher Scientific #25200072), combined in groups of 4 wells per tube, counted, centrifuged at 300 x g for 10 minutes, and resuspended in 100% FBS to a density of 1.0-1.2 × 10^6^ cells/mL. To freeze, cells in FBS were mixed in a 1:1 volume with FBS + 20% DMSO (final concentration 5-6 × 10^5^ cells in 10% DMSO) and frozen in 1 mL aliquots at −80°C overnight in a Mr. Frosty cryo freezing container before transferring to liquid nitrogen.

### T4-AVV preparation

#### Protein and mRNA display

GG-Hoc was added to purified heads in a 20:1 molar ratio (GG-Hoc: Hoc binding sites) and incubated at room temperature (RT) for 1 hour, rotating. After GG-Hoc binding, Soc fusion proteins were added at 10:1 molar ratio (Soc-fusion: Soc binding sites) and incubated at RT for 1 hour rotating. The protein decorated heads were pelleted at 30,000 x g for 1 hour at 4°C and unbound GG-Hoc and Soc-fusion proteins were removed from the supernatant. Head pellets were washed twice with 1 mL 1X PBS (pH 7.4) with a centrifugation step (30,000 x g, 5 min, 4°C) after each wash and resuspended in 1X PBS for further biomolecule binding. To display targeting antibodies, protein decorated heads were mixed with mAbs at 2.5:1 molar ratio of mAb to Hoc binding sites and incubated at RT for 1 hour rotating. Heads were centrifuged and washed as before to remove unbound mAb. Prior to mRNA display, protein coated T4-AVV could be stored at 4°C for 24-72 hours. Finally, mRNA was displayed on the surface by mixing protein and mAb decorated heads in a 0.5:1 molar ratio of mRNA to Soc binding sites and incubated at 8°C for 1 h followed by centrifugation and washing as before to remove unbound mRNA. Completed, mRNA-coated T4-AVV were resuspended in sterile 1X PBS pH 7.4 and used immediately for lipid coating and transduction.

#### Lipid coating

Prepared T4-AVVs were diluted to the final MOI in sterile 1X PBS pH 7.4 and 0.25-0.5 µL of cationic lipid transfection reagent was added such that the final volume of components totaled 20 µL per well. For mRNA-LNP controls, 100 ng of reporter mRNA was diluted with sterile 1X PBS pH 7.4 and lipid transfection reagent to a final volume of 20 µL.. All tubes were vortexed for 5s and incubated at RT for 15 minutes. After incubation, 80 µL of Stemspan SFEM II supplemented with 1X CD34 expansion supplement, 1 µM UM729 and PS was added to each sample to form the complete transduction mixture.

### Protein and mRNA display analysis

Prior to lipid coating, aliquots of T4-AVVs were saved for quantification analysis. To quantify protein display, T4-AVVs were mixed with Laemmli SDS sample buffer (ThermoFisher Scientific J61337.AD), boiled for 5 minutes and proteins were separated by SDS-PAGE and stained by Coomassie R-250 (Biorad#1610436). The density of protein bands was measured by ImageJ and compared to the density of gp23 or a BSA standard of known concentration to estimate copy numbers. To quantify mRNA display, T4-AVVs were mixed with a digestion cocktail containing equal volumes of Proteinase K (ThermoFisher Scientific #EO0491), 0.5 M EDTA, pH 8.0 (Quality Biological #351-027-721) and 10% SDS (Quality Biological #351-032-101) and incubated at 65°C for 30 minutes. The digested T4-AVV were mixed with purple loading dye (NEB #B7024S) and separated on 1% (wt/vol) agarose gel electrophoresis. After visualization, the density of mRNA bands was measured by ImageJ and compared to the density mRNA standards of known concentration to estimate copy number.

### HSC transduction

One day prior to transduction, one vial of pre-expanded HSCs was thawed as previously described, resuspended in expansion media and counted. Cells were plated in 96-well plates at 15,000 cells per well in 100 µL media and incubated 24 hours at 37°C, 5% CO_2_. To transduce, 80 µL of media was removed from then cells and replaced with 100 µL of the prepared transduction mixture. The cells were incubated at 37°C, 5% CO_2_ for the time listed in each experiment.

### Cell viability assays

To count cell numbers, the supernatant was saved from each well and pelleted at 500 x g for 1 minute. Adherent cells were removed by the addition of 50 µL 0.25% Trypsin with incubation for 5 minutes at 37°C, 5% CO2. Trypsin was quenched with 50 µL HSC expansion medium and the detached cells were combined with the supernatant pellet. Cells were pipetted up and down gently to achieve a homogenous cell suspension and 20 µL of suspension was mixed with 20 µL 0.4% Trypan blue. Cells counts and viability were determined by the QuadCount Automated Cell Counter (Accuris Instruments). Cell viability was also assessed using Cell Titer-Glo (Promega #G7571) per manufacturer’s instructions. After room temperature equilibration, 100 µL of Cell Titer-Glo reagent was added to each well containing 100 µl of media, cells were lysed by orbital shaking for 2 minutes and incubated at RT for 10 minutes. After incubation the 200 µL sample was transferred to opaque 96-well plates and luminescence was read on the Glomax Multi Detection System (Promega) with 0.5 s integration time.

### Microscopy

To assess viability, cells were stained with 0.2% trypan blue and incubated at RT for 5 min. The media containing trypan blue was removed, cells were washed 1x in 100 µL PBS and resuspended in 100 µL PBS. Cells were imaged immediately in the well using an inverted microscope (Axio Observer, Carl Zeiss) to observe cells that had internalized the dye. To measure GFP expression, at the indicated time points post transduction, plates were removed from the incubator and imaged directly in the well on an inverted microscope (Axio Observer, Carl Zeiss) using brightfield and fluorescence modes, To quantify transduction efficiency, the number of GFP positive cells per field was counted and divided by the total number of cells in that field. Two to three fields of at least 100 total cells were counted per condition.

### Luminescence measurement

Luminescence was measured by modifying manufacturer’s instructions (Promega #E1501). The supernatant from each well was saved and centrifuged at 1500 rpm for 5 minutes to collect any nonadherent cells. Twenty microliters of 1X lysis buffer was added to the cell pellet and to each sample well to lyse remaining adherent cells. Cells were incubated at room temperature for 5-10 minutes to lyse. The 40 µL lysis buffer was combined, vortexed and centrifuged to pellet debris. Then 20 µL of supernatant was transferred to a 96-well opaque white plate and 100 µL luciferase assay reagent was added to each well. Luminescence was read immediately on the Glomax Multi Detection System (Promega) with 0.5 s integration time. Luminescence assays were performed in biological duplicate or triplicate.

### β-galactosidase activity measurement

The conversion of ONPG was used to measure Soc-β-gal activity per manufacturer’s instructions (Promega #E2000). The supernatant from each well was saved, the wells were washed with 100 µL PBS and the wash volume was combined with the supernatant. These nonadherent cells were pelleted by centrifugation at 1500 rpm for 5 minutes. The cell pellet was lysed with 37 µL reporter lysis buffer and adherent cells in the well were lysed with the same volume. Cells were allowed to lyse for 10-15 minutes and lysates from adherent and nonadherent fractions or each well were combined. To measure Soc-β-gal activity 50 µL of each lysate was transferred to a clear 96-well plate assay plate, mixed with 50 µL 2X assay buffer and incubated at 37°C for 30 minutes covered. The reactions were stopped by the addition of 150 µL 1 M sodium carbonate and the absorbance at 420 nm was read immediately on a SpectraMax 340PC384 microplate reader (Molecular Devices). Purified Soc-β-gal (13 ng diluted in gel filtration buffer) was used as a positive control for ONPG conversion. Soc-β-gal activity measurements were performed in biological duplicate or triplicate.

## Acknowledgements

This research was supported by NIDA, NIH Avant Garde Award 1DP1DA060580-01 to V.B.R

## Author Contributions

VBR designed and directed the project. CES, JZ and VBR performed experimental design. CES, CDV, OD, and KT produced biological reagents. CES performed experiments and analyzed data. CES and VBR wrote the manuscript.

