## Supplemental Information for "Targeted Delivery of Nucleic Acid and Protein Cargos into Primary Human Hematopoietic Stem Cells Using Bacteriophage T4"

**Supplementary Information**

### Supplemental Figure 1

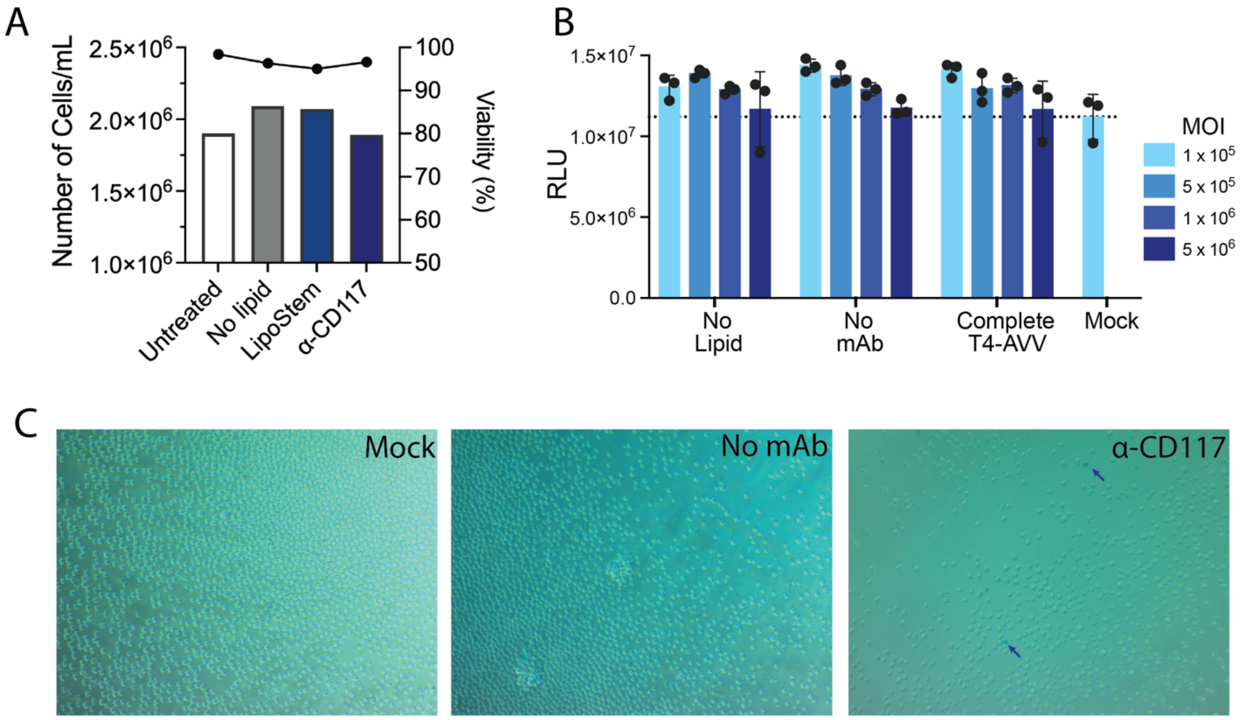


Supplemental Figure 1. T4-AVV transduction does not affect HSC viability. **A.** Total cell count (bars) and percent viability (line) of HSC 48 hpt with the indicated T4-AVV variations. Cells were removed from the well with trypsin, mixed with 0.2% trypan blue and analyzed on a Quadcount automated counter. **B**. Cells were transduced in triplicate by the indicated T4-AVVs at increasing MOIs and cell viability was analyzed by Cell Titer Glo assay. **C**.  Representative bright field images of duplicate wells in **A**. Cells were stained with 0.2% trypan blue for 5 min, washed and visualized for uptake of the dye. Arrows indicate cells that have taken up the stain. Data for **A-C** are representative of two independent experiments.

### Supplemental Note 1. DNA templates and recombinant proteins.

#### *In vitro* transcription templates

#### Codon-optimized luciferase, CDS green

AGTAATACGACTCACTATAAGGAGAACTCTTCTGGTCCCCACAGACTCAGAGAGAACCCACCAT
GGAAGATGCCAAAAACATTAAGAAGGGCCCAGCGCCATTTTACCCACTCGAAGACGGGACCGCC
GGCGAGCAGCTGCACAAAGCCATGAAGAGGTACGCCCTGGTGCCCGGCACCATCGCCTTTACCG
ACGCACATATCGAGGTGGACATTACCTACGCCGAGTACTTCGAGATGAGCGTTAGGCTGGCAGA
AGCTATGAAGAGGTATGGGCTGAATACAAACCATAGGATCGTGGTGTGCAGCGAGAATAGCCTG
CAGTTCTTCATGCCCGTGCTGGGCGCCCTGTTCATCGGCGTGGCTGTGGCCCCAGCTAACGACA
TCTACAACGAGAGGGAGCTGCTGAACAGCATGGGCATCAGCCAGCCCACCGTCGTATTCGTGAG
CAAGAAAGGGCTGCAGAAGATCCTCAACGTGCAGAAGAAGCTCCCCATCATACAGAAGATCATC
ATAATGGATAGCAAGACCGACTACCAGGGCTTCCAGAGCATGTACACCTTCGTGACTTCCCATC
TGCCACCCGGCTTCAACGAGTACGACTTCGTGCCCGAGAGCTTCGACAGGGACAAAACCATCGC
CCTGATCATGAACAGTAGTGGCAGTACCGGACTGCCCAAGGGCGTAGCCCTTCCCCACAGGACC
GCTTGTGTCCGATTCAGTCATGCCAGGGACCCCATCTTCGGCAACCAGATCATCCCCGACACCG
CTATCCTCAGCGTGGTGCCATTTCACCACGGCTTCGGCATGTTCACCACGCTGGGCTACCTGAT
CTGCGGCTTTAGGGTCGTGCTCATGTACAGGTTCGAGGAGGAGCTGTTCCTGAGGAGCCTGCAG
GACTATAAGATTCAGTCTGCCCTGCTGGTGCCCACACTGTTTAGCTTCTTCGCTAAGAGCACTC
TCATCGATAAGTACGACCTCAGCAACCTGCACGAGATCGCCAGCGGCGGGGCGCCCCTCAGCAA
GGAGGTAGGCGAGGCCGTGGCCAAAAGGTTCCACCTGCCAGGCATCAGGCAGGGCTACGGCCTG
ACAGAAACAACCAGCGCCATTCTGATCACCCCCGAAGGGGACGACAAGCCTGGCGCAGTAGGCA
AGGTGGTGCCCTTCTTCGAGGCTAAGGTGGTGGACCTGGACACCGGCAAGACACTGGGCGTGAA
CCAGAGGGGCGAGCTGTGCGTCAGGGGCCCCATGATCATGAGCGGCTACGTTAACAACCCCGAG
GCTACAAACGCTCTCATCGACAAGGACGGCTGGCTGCACAGCGGCGACATCGCCTACTGGGACG
AGGACGAGCACTTCTTCATCGTGGACAGGCTGAAGAGCCTGATCAAATACAAGGGCTACCAGGT
AGCCCCAGCCGAACTGGAGAGCATCCTGCTGCAGCACCCCAACATCTTCGATGCCGGGGTCGCT
GGCCTGCCCGACGACGATGCTGGGGAACTGCCCGCCGCAGTCGTCGTGCTGGAACACGGCAAAA
CCATGACCGAGAAGGAGATCGTGGACTATGTGGCCAGCCAGGTTACAACCGCCAAGAAGCTGAG
GGGCGGCGTTGTGTTCGTGGACGAGGTGCCTAAAGGACTGACCGGCAAGCTGGACGCCAGGAAG
ATCAGGGAGATTCTCATTAAGGCCAAGAAGGGCGGCAAGATCGCCGTGTGAGCTGGAGCCTCGG
TGGCCATGCTTCTTGCCCCTTGGGCCTCCCCCCAGCCCCTCCTCCCCTTCCTGCACCCGTACCC
CCGTGGTCTTTGAATAAAGTCTGAGTGGGCGGCAAAAAAAAAAAAA

#### EGFP CDS green

AGTAATACGACTCACTATAAGGAGAACTCTTCTGGTCCCCACAGACTCAGAGAGAACCCACCAT
GGTGAGCAAGGGCGAGGAGCTGTTCACCGGGGTGGTGCCCATCCTGGTCGAGCTGGACGGCGAC
GTAAACGGCCACAAGTTCAGCGTGTCCGGCGAGGGCGAGGGCGATGCCACCTACGGCAAGCTGA
CCCTGAAGTTCATCTGCACCACCGGCAAGCTGCCCGTGCCCTGGCCCACCCTCGTGACCACCCT
GACCTACGGCGTGCAGTGCTTCAGCCGCTACCCCGACCACATGAAGCAGCACGACTTCTTCAAG

TCCGCCATGCCCGAAGGCTACGTCCAGGAGCGCACCATCTTCTTCAAGGACGACGGCAACTACA
AGACCCGCGCCGAGGTGAAGTTCGAGGGCGACACCCTGGTGAACCGCATCGAGCTGAAGGGCAT
CGACTTCAAGGAGGACGGCAACATCCTGGGGCACAAGCTGGAGTACAACTACAACAGCCACAAC
GTCTATATCATGGCCGACAAGCAGAAGAACGGCATCAAGGTGAACTTCAAGATCCGCCACAACA
TCGAGGACGGCAGCGTGCAGCTCGCCGACCACTACCAGCAGAACACCCCCATCGGCGACGGCCC
CGTGCTGCTGCCCGACAACCACTACCTGAGCACCCAGTCCGCCCTGAGCAAAGACCCCAACGAG
AAGCGCGATCACATGGTCCTGCTGGAGTTCGTGACCGCCGCCGGGATCACTCTCGGCATGGACG
AGCTGTACAAGTAAGCTGGAGCCTCGGTGGCCATGCTTCTTGCCCCTTGGGCCTCCCCCCAGCC
CCTCCTCCCCTTCCTGCACCCGTACCCCCGTGGTCTTTGAATAAAGTCTGAGTGGGCGGCAAAA
AAAAAAAA

**Recombinant proteins**

**GG-Hoc**

GSSHHHHHHSSGLVPRGSHMASSLVTGSMGTPAVTTYKLVINGKTLKGETTTKAVDAE
TAEKAFKQYANDNGVDGVWTYDDATKTFTVTEVNTPAVTTYKLVINGKTLKGETTTKAV
DAETAEKAFKQYANDNGVDGVWTYDDATKTFTVTELEFTVDITPKTPTGVIDETKQFTA
TPSGQTGGGTITYAWSVDNVPQDGAEATFSYVLKGPAGQKTIKVVATNTLSEGGPETA
EATTTITVKNKTQTTTLAVTPASPAAGVIGTPVQFTAALASQPDGASATYQWYVDDSQV
GGETNSTFSYTPTTSGVKRIKCVAQVTATDYDALSVTSNEVSLTVNKKTMNPQVTLTPP
SINVQQDASATFTANVTGAPEEAQITYSWKKDSSPVEGSTNVYTVDTSSVGSQTIEVTA
TVTAADYNPVTVTKTGNVTVTAKVAPEPEGELPYVHPLPHRSSAYIWCGWWVMDEIQK
MTEEGKDWKTDDPDSKYYLHRYTLQKMMKDYPEVDVQESRNGYIIHKTALETGIIYTYP

**Cas9-Soc**

PKKKRKVMDKKYSIGLDIGTNSVGWAVITDEYKVPSKKFKVLGNTDRHSIKKNLIGALLF
DSGETAEATRLKRTARRRYTRRKNRICYLQEIFSNEMAKVDDSFFHRLEESFLVEEDKK
HERHPIFGNIVDEVAYHEKYPTIHLRKKLVDSTDKADLRLIYLALAHMIKFRGHFLIEGDL
NPDNSDVDKLFIQLVQTYNQLFEENPINASGVDAKAILSARLSKSRRLENLIAQLPGEKK
NGLFGNLIALSLGLTPNFKSNFDLAEDAKLQLSKDTYDDDLDNLLAQIGDQYADLFLAAK
NLSDAILLSDILRVNTEITKAPLSASMIKRYDEHHQDLTLLKALVRQQLPEKYKEIFFDQS
KNGYAGYIDGGASQEEFYKFIKPILEKMDGTEELLVKLNREDLLRKQRTFDNGSIPHQIH
LGELHAILRRQEDFYPFLKDNREKIEKILTFRIPYYVGPLARGNSRFAWMTRKSEETITP
WNFEEVVDKGASAQSFIERMTNFDKNLPNEKVLPKHSLLYEYFTVYNELTKVKYVTEG
MRKPAFLSGEQKKAIVDLLFKTNRKVTVKQLKEDYFKKIECFDSVEISGVEDRFNASLG
TYHDLLKIIKDKDFLDNEENEDILEDIVLTLTLFEDREMIEERLKTYAHLFDDKVMKQLKR
RRYTGWGRLSRKLINGIRDKQSGKTILDFLKSDGFANRNFMQLIHDDSLTFKEDIQKAQ
VSGQGDSLHEHIANLAGSPAIKKGILQTVKVVDELVKVMGRHKPENIVIEMARENQTTQ
KGQKNSRERMKRIEEGIKELGSQILKEHPVENTQLQNEKLYLYYLQNGRDMYVDQELDI
NRLSDYDVDHIVPQSFLKDDSIDNKVLTRSDKNRGKSDNVPSEEVVKKMKNYWRQLLN
AKLITQRKFDNLTKAERGGLSELDKAGFIKRQLVETRQITKHVAQILDSRMNTKYDENDK
LIREVKVITLKSKLVSDFRKDFQFYKVREINNYHHAHDAYLNAVVGTALIKKYPKLESEFV

YGDYKVYDVRKMIAKSEQEIGKATAKYFFYSNIMNFFKTEITLANGEIRKRPLIETNGETG
EIVWDKGRDFATVRKVLSMPQVNIVKKTEVQTGGFSKESILPKRNSDKLIARKKDWDPK
KYGGFDSPTVAYSVLVVAKVEKGKSKKLKSVKELLGITIMERSSFEKNPIDFLEAKGYKE
VKKDLIIKLPKYSLFELENGRKRMLASAGELQKGNELALPSKYVNFLYLASHYEKLKGSP
EDNEQKQLFVEQHKHYLDEIIEQISEFSKRVILADANLDKVLSAYNKHRDKPIREQAENII
HLFTLTNLGAPAAFKYFDTTIDRKRYTSTKEVLDATLIHQSITGLYERIDLSQLGGDGGG
GSRSGGYVNIKTFTHPAGEGKEVKGMEVSVPFEIYSNEHRIADAHYQTFPSEKAAYTV
VTDAADWRTKNAAMFTPTPVSGHHHHHH

**Soc-GG-TAT**

GSSHHHHHHSSGASTRGYVNIKTFEQKLDGNKKIEGKEISVAFPLYSDVHKISGAHYQT
FPSEKAAYSTVYEENQRTEWIAANEDLWKVTGSSGLVPRGSHMASSLVTGSMGTPAVT
TYKLVINGKTLKGETTTKAVDAETAEKAFKQYANDNGVDGVWTYDDATKTFTVTEVNTP
AVTTYKLVINGKTLKGETTTKAVDAETAEKAFKQYANDNGVDGVWTYDDATKTFTVTEL
EGGGSGGSGGSGGSLVPRGSHMASNGYGRKKRRQRRR

**Soc-β-galactosidase**

GSSHHHHHHSSGASTRGYVNIKTFEQKLDGNKKIEGKEISVAFPLYSDVHKISGAHYQT
FPSEKAAYSTVYEENQRTEWIAANEDLWKVTGSSGMTMITDSLAVVLQRRDWENPGV
TQLNRLAAHPPFASWRNSEEARTDRPSQQLRSLNGEWRFAWFPAPEAVPESWLECD
LPEADTVVVPSNWQMHGYDAPIYTNVTYPITVNPPFVPTENPTGCYSLTFNVDESWLQ
EGQTRIIFDGVNSAFHLWCNGRWVGYGQDSRLPSEFDLSAFLRAGENRLAVMVLRWS
DGSYLEDQDMWRMSGIFRDVSLLHKPTTQISDFHVATRFNDDFSRAVLEAEVQMCGE
LRDYLRVTVSLWQGETQVASGTAPFGGEIIDERGGYADRVTLRLNVENPKLWSAEIPNL
YRAVVELHTADGTLIEAEACDVGFREVRIENGLLLLNGKPLLIRGVNRHEHHPLHGQVM
DEQTMVQDILLMKQNNFNAVRCSHYPNHPLWYTLCDRYGLYVVDEANIETHGMVPMN
RLTDDPRWLPAMSERVTRMVQRDRNHPSVIIWSLGNESGHGANHDALYRWIKSVDPS
RPVQYEGGGADTTATDIICPMYARVDEDQPFPAVPKWSIKKWLSLPGETRPLILCEYAH
AMGNSLGGFAKYWQAFRQYPRLQGGFVWDWVDQSLIKYDENGNPWSAYGGDFGDT
PNDRQFCMNGLVFADRTPHPALTEAKHQQQFFQFRLSGQTIEVTSEYLFRHSDNELLH
WMVALDGKPLASGEVPLDVAPQGKQLIELPELPQPESAGQLWLTVRVVQPNATAWSE
AGHISAWQQWRLAENLSVTLPAASHAIPHLTTSEMDFCIELGNKRWQFNRQSGFLSQM
WIGDKKQLLTPLRDQFTRAPLDNDIGVSEATRIDPNAWVERWKAAGHYQAEAALLQCT
ADTLADAVLITTAHAWQHQGKTLFISRKTYRIDGSGQMAITVDVEVASDTPHPARIGLNC
QLAQVAERVNWLGLGPQENYPDRLTAACFDRWDLPLSDMYTPYVFPSENGLRCGTRE
LNYGPHQWRGDFQFNISRYSQQQLMETSHRHLLHAEEGTWLNIDGFHMGIGGDDSW
SPSVSAEFQLSAGRYHYQLVWCQK
